# Impaired renal base excretion in secretin receptor knock-out mice during prolonged base-loading

**DOI:** 10.64898/2026.03.05.709818

**Authors:** Tobias Jensen, Jesper Frank Andersen, Laura Woidemann Trans, Ida Marie Modvig, Helga Vitzthum, Jens Juul Holst, Bolette Hartman, Samuel L Svendsen, Mads Vaarby Sørensen, Jens Leipziger, Peder Berg

## Abstract

**Aim:** Secretin was recently found to play a pivotal role in the renal adaptation to acute base excess. Here, secretin increases pendrin-dependent HCO_3_^-^ secretion from the beta-intercalated cells in the cortical collecting ducts. Whether secretin and its receptor play a role during prolonged base-loading remains unknown.

**Methods:** Urine and blood acid-base analyses were carried out in secretin receptor (SCTR) KO and WT mice at baseline and after 1 and up to 8 days of base-loading with NaHCO_3_-enriched drinking water. Changes in pendrin protein abundance and function were assessed by immunoblotting and isolated tubule perfusion experiments. Plasma secretin levels and renal SCTR expression were assessed after 24 hours of acid/base-loading by radioimmunoassay and qPCR, respectively.

**Results:** SCTR KO mice responded with diminished urine alkalization and a lesser reduction of urinary acid excretion when base-loaded for 48 hours. Concordantly, SCTR KO mice presented with increased blood base retention compared with WTs. Base-loaded SCTR WT and KO mice showed comparable total pendrin protein abundance. Despite this, pendrin function was markedly lower in SCTR KO mice. Base-loaded mice had higher plasma secretin and renal SCTR levels compared with acid-loaded mice. Higher arterial HCO_3_ ^-^ associated with higher renal SCTR mRNA expression.

**Conclusion:** Loss of the SCTR diminishes renal base excretion capacity and exacerbates systemic base accumulation during prolonged base-loading. Further, plasma secretin and renal SCTR mRNA levels are modulated by acid-base intake. These findings further support a central role of secretin and its receptor in the regulation of both acute and prolonged base excess.

**Key points:**

1. In mice, the receptor of the gut hormone secretin is required for the kidney to adequately handle a sustained base load. In mice with secretin receptor knockout, renal base excretion is impaired and metabolic alkalosis is aggravated and slower to resolve.
2. Base-loaded knockout mice up-regulate pendrin protein abundance normally, the bicarbonate-secreting Cl^-^/HCO_3_^-^exchanger in collecting-duct β-intercalated cells, but exhibit lower pendrin function. This suggests that the defect lies in the activation of pendrin, rather than in the regulation of its abundance.
3. Renal secretin-receptor expression increases with higher arterial bicarbonate and plasma secretin, and renal secretin receptor expression, levels are lower in acid-loaded mice compared to base-loaded mice. This indicates that the secretin/secretin receptor system is itself affected by acid-base status/acid-base intake.

## Introduction

Secretin was discovered by Bayliss and Starling in 1902,^1^ creating the foundation of our modern understanding of hormone physiology. Beyond its known roles in gastro-intestinal physiology, secretin has been demonstrated to elicit a variety of effects such as systemic vasodilation and increased cardiac output,^2–4^ modulation of vasopressin release,^5,6^ regulation of thirst,^7^ CNS-mediated reduction of appetite and increased satiety, along with activation of brown adipose tissue.^8–10^ Secretin also governs epithelial HCO_3_^-^ secretion in a variety of organs, including the pancreatic ducts,^11^ hepatic bile ducts^12^ and collecting ducts of the kidney,^13–15^ the latter recently described by our research group.^13,14,16^ Renal HCO_3_ ^-^ secretion is mediated through the Cl^-^/HCO_3_ ^-^-exchanger pendrin (SLC26A4) positioned at the apical membrane of the beta-intercalated cells (beta-ICs) in the collecting duct.^17^ The regulation and function of pendrin is critically dependent on the CFTR anion channel, also located at the apical membrane.^13,18,19^ Loss of either pendrin or CFTR impairs renal base excretion, aggravating both acute and chronic metabolic alkalosis in mice and humans,^13,19–23^ which eventually triggers partial respiratory compensation.^20^

Secretin elicits its renal effects via the secretin receptor (SCTR) expressed on the basolateral membrane of the beta-ICs in the connecting tubule and cortical collecting duct.^13,14,24,25^ SCTR activation acutely increases pendrin function and hence urinary HCO_3_^-^ excretion.^13,14,26,27^ Acute metabolic alkalosis elevates plasma secretin levels, supporting the role of secretin in the regulation of systemic HCO_3_ ^-^ levels.^13^ In accordance with this, mimicking metabolic alkalosis in isolated perfused upper small intestines increases the release of secretin as measured in portal venous blood.^14^

Underlining the importance of a functional response to secretin, it was recently demonstrated that SCTR KO mice present with strongly impaired renal base excretion capabilities during acute base-loading. Consequently, SCTR KO mice exhibit aggravated metabolic alkalosis and compensatory hypoventilation under these circumstances.^14^

The main purpose of this study was to investigate the renal response to prolonged base-loading in SCTR KO mice. We hypothesized that SCTR KO mice would present with decreased urinary base excretion and augmented elevation of plasma HCO_3_^-^ during prolonged base-loading. Furthermore, we hypothesized that plasma secretin levels and SCTR expression in the kidney could be affected by systemic acid-base status.

To examine this, a variety of methods were employed, i.e., metabolic cage studies, systemic and urine acid-base measurements, immunoblotting, perfusion of isolated cortical collecting ducts, semi-quantitative PCR of SCTR in kidney tissue and radioimmunoassay of secretin in plasma.

We found that prolonged base-loading disturbed blood acid-base parameters more in SCTR KO mice compared with WT mice. Concordantly, SCTR KO mice exhibited diminished urine alkalization capabilities. Pendrin activity following base-loading was strongly diminished in SCTR KO mice despite similar protein abundance between KOs and WTs. Lastly, base-loaded mice had higher plasma secretin levels and renal SCTR expression than acid-loaded animals and a higher arterial HCO_3_^-^ was associated with increased renal SCTR expression after 24 hours of acid/base-loading.

These results offer a comprehensive understanding of secretin’s role in renal acid-base handling and suggest that a functional secretin response is necessary to appropriately adapt to a prolonged alkali load.

## Methods

### Animals

Experiments and handling of mice were approved by the Danish animal welfare regulations (2016-15-0201-01129 and 2022-15-0201-01129). At Aarhus University animals are only used in accordance with applicable legislation, ethical standards and guidelines set by national and European authorities. Mice were housed with a 12-hour day/night cycle and *ad libitum* access to drinking water and standard rodent chow. Experiments including genetically modified mice were bred from heterozygous breeding pairs.

SCTR KO mice were generated as previously explained and were in the C57Bl/6J genetic background.^5, 14^ Mice of both sexes of similar age (>7 weeks) were used for *in vivo* experiments. Mice of 4-8 weeks were used for tubule perfusion experiments. Male and female C57Bl/6J mice, 50% female, 10-12 weeks of age, were purchased from Janvier. These were allowed at least 1 week of acclimatization before entering experiments.

### Acid/base-loading

During base-loading, standard drinking water was substituted with *ad libitum* MilliQ water with either 112 or 160 mM NaHCO_3_. KOs were exposed to 112 mM NaHCO_3_ drinking water and WTs were given either 112 or 160 mM NaHCO_3_ drinking water, the latter in an attempt to adjust for the previously reported ∼40% higher water intake in SCTR KO mice that otherwise could lead to a higher total intake of NaHCO_3_ in KO mice.^5^ In our lab, a similarly elevated water intake was observed with 112 mM NaHCO_3_ drinking water (Figure S1). The bicarbonate intake itself was not measured directly in all experiments and, as such, the exposure between genotypes should therefore be regarded as approximate rather than exactly matched.

C57Bl/6J mice were acid/base-loaded by changing the drinking water to MilliQ with either 200 mM NaHCO_3_ or 200 mM NH_4_Cl for 24 hours ahead of plasma and kidney tissue sampling.

### Urine collection and measurement of acid-base parameters

Urine was collected under mineral oil in metabolic cages every 24 hours. Urine pH was measured with a micro pH electrode (pH-500, Unisense, Aarhus, Denmark). Ammonium was measured using an Orion High-performance Ammonia Ion-selective Electrode (Thermo Scientific, cat. No. 9512HPBNWP), with the use of an Ammonia pH-adjusting Ionic Strength Adjuster (Thermo Scientific, Cat. No. 951211) as previously described.^28^ Titratable acid (TA) was measured by titration using the method of Chan^29^ modified for small-volume samples.^28^

### Blood gas analysis

Venous tail blood was sampled from lightly anesthetized mice (isoflurane anesthesia). Venous blood samples were drawn during baseline conditions and after 1, 4 and 8 days of base-loading. During mechanical normal ventilation, arterial blood samples were taken from the tail artery of ketamine-xylazine anaesthetized mice. Blood was sampled with glass capillaries (CLINITUBES, Radiometer, Denmark) and analyzed with an ABL80 Flex Blood gas analyzer (Radiometer, Denmark) immediately after collection. Comprehensive blood acid base analysis is given by the automated blood gas analyzer including PaCO_2_, pH and calculated standard HCO_3_ ^-^.

### Measurement of pendrin activity in isolated perfused collecting ducts

Measurement of pendrin activity was performed as previously described.^13,14^ In short, cortical collecting ducts were manually dissected and perfused with a pipette system at 37°C. A pH-sensitive dye BCECF-AM (2’,7’-bis-(2-carboxyethyl)-5-(and-6)-Carboxyfluorescein, Acetoxymethyl Ester) was loaded from the luminal side, where it primarily loads into intercalated cells.^30^ Single cell intracellular pH was then monitored throughout the experiments, and pendrin activity was determined as the initial alkalization rate upon fast luminal chloride removal.^31^ The analyzing person who calculated the initial alkalization rates was blinded to animal genotype. Pendrin-positive cells can be unequivocally identified and discerned from type-A ICs by fast luminal chloride removal. Only pendrin positive cells respond with an immediate intracellular alkalization.^31^

Intrinsic intracellular buffer capacity (β_i_) was estimated from intracellular pH using previous data from our lab^32^ showing a very high correlation between intracellular pH and intrinsic buffer capacity (Pearsons’s correlation coefficient: 0.90, Figure S2). In the presence of CO_2_/HCO_3_ ^-^, the total buffer capacity (β) is equal to β_i_+2.3×(HCO ^-^). Intracellular HCO ^-^ concentration was calculated as: [HCO ^-^] =0.03 x PCO x 10^pHi-6.1^, with the assumption that intracellular PCO_2_ equals extracellular PCO_2_ (37 mm Hg in solutions gassed with 5% CO_2_ and 95% O_2_). Base flux was estimated as dpH_i_/dt x β_total_ (in mM/min).

### Mechanical ventilation of C57Bl/6J mice, tissue harvesting and blood sampling

During ketamine-xylazine anaesthesia, mice were positioned on an intubation platform (Kent Scientific endotracheal intubation kit) and intubated with a 20G venflon tube equipped with a safety wedge, using a wire guide to secure placement into the trachea. Mechanical ventilation was provided with a Minivent mouse ventilator (Model 845, Hugo Sachs Elektronik – Harvard Apparatus) and continuously monitored with capnography (Capnograph Type 340, Hugo Sachs Elektronik – Harvard Apparatus).

Normal ventilation was achieved by adjusting ventilatory parameters according to an end-tidal CO_2_ of approximately 2.9% and was confirmed by an arterial pCO2 of ∼40 mmHg (from arterial blood gas analysis as described above).

During normal ventilation, blood was drawn from the left ventricle and immediately transferred to a heparin-coated tube (BD microtainer tubes, cat nr. 365986) and put on ice. Plasma was separated by centrifugation (9000 rpm for 2 minutes at 4°C). A full kidney was harvested and snap frozen in liquid nitrogen for subsequent qPCR analysis and immunoblotting.

### Immunoblotting

Tissue harvesting for Western blots was performed as previously described.^33^ In short, blotting was performed on the supernatant of halved kidneys processed in lysis buffer (sucrose: 0.3 M, imidazole: 25 mM, leupeptin: 8.5 mM, Pefabloc (Sigma): 1 mM, and phosSTOP (Roche Diagnostics, Manheim, Germany): 1 tablet/10 ml) with a tissuelyser (Qiagen) for 30 seconds and spun for 15 minutes (1000 g).

Protein concentrations were measured using Pierce^TM^ BCA Protein Assay Kit. All samples were run on Criterion TGX Precast Gels (Bio-Rad). 10 µg of total protein was loaded into each sample well. This was validated to be within the linear detection range for the used pendrin antibody.^19^ The pendrin antibody was diluted 1:1000. Membranes were developed using the Clarity Western enhanced Chemiluminescence substrate (Bio-Rad) in an ImageQuant LAS 4000 mini (GE Healthcare Life Science). Density signals of pendrin were normalized to the signal of a Coomassie staining of the membranes post chemiluminescent detection of the protein of interest to enable correction for differences in protein loading.^34^ The pendrin antibody was a kind gift from Associate Professor Sebastian Frische and was previously described.^35^

### Radioimmune assay of plasma secretin

Plasma secretin levels were measured using a previously developed and validated radioimmunoassay using a C-terminally directed secretin antibody (codename 5595-3) kindly provided by Professor Jan Fahrenkrug (Department of Clinical Biochemistry, Bispebjerg Hospital, Copenhagen, Denmark).^36^ Mouse secretin and radioactive labeled mouse secretin were obtained from Phoenix Pharmaceuticals, Inc (cat. no. 067-04 and T-067-06, CA). Three samples were >1.5 IQR above the 75^th^ percentile of the full cohort and were excluded from statistical analysis.

### Semi-quantitative PCR of kidney tissue

Initially, half a kidney was homogenised in Qiazol Lysis Reagent (Cat. No. 79306, QIAGEN) using Precellys 24. The remaining half was saved for backup. RNA was then purified using the RNeasy Mini Kit (Cat. No. 74104, QIAGEN) and cDNA was generated with FastGene Scriptase II cDNA Synthesis Kit LS63 (NIPPON Genetics EUROPE). Real-time PCR was then performed with RealQ 2x Master Mix Green Without ROX^TM^ (Cat. No. A323406 AMPLIQON IIII) against the SCTR (primers: forward, TCAAGGACGCCGTACTCTTC; reverse, CGAAGAGCACAAATGCCTGC).

Housekeeping genes: mPPIA (primers: forward, GTGGTCTTTGGGAAGGTG AA; reverse, TTACAGGACATTGCGAGCAG) and mHPRT (forward, AAGCTTGCTGGTGAAAAGGA; reverse, TTGCGCTCATCTTAGGCTTT).

SCTR expression was normalized to the geometric mean of the housekeeping genes and differences between experimental groups were assessed using the delta-delta Cq method with to the control group as reference.

### Statistics

Data distribution was assessed by inspection of QQ-plots. If necessary, data were log transformed before analysis. Differences between two groups were assessed using a student’s t-test. Differences between more than two groups were assessed using one-way ANOVA. Differences between two groups with time-dependent data or more than one intervention were assessed using two-way ANOVA or mixed-effects analyses where appropriate. Genotype specific difference between slopes and intercepts for the association between blood bicarbonate and acid excretion and between intracellular pH and pendrin function were assessed by inclusion of a bicarbonate-by-genotype/pH_i_-by-genotype interaction term in the regression analysis. If the interaction term was significant (F-test), it was concluded that genotype had a significant effect modification on the assessed association. If the interaction term was not significant, it was removed and the intercept (the effect of genotype after adjustment for pH_i_) was assessed using a common slope for both genotypes (F-test). All tests were two-tailed and performed at a significance level of 5%. Each figure legend explains what tests were used to assess statistical significance.

## Results

### SCTR KO mice respond with diminished urine alkalization during 48 hours of base-loading

To investigate the role of the SCTR during prolonged base excess, a series of metabolic cage experiments were performed on SCTR WT and KO mice. Urine was collected after 24 and 48 hours with standard drinking water. Following this, the drinking water was changed to NaHCO_3_-enriched water for 48 hours. KO mice received 112 mM NaHCO_3_, WT mice received 112 mM NaHCO_3_ or 160 mM NaHCO_3_ (160 mM was chosen to adjust for the ∼40% higher water intake in SCTR KO mice, see Chu et al^5^ and Figure S1). Urine was collected again after 24 and 48 hours of base-loading.

In congruence with our previous findings,^14^ no differences in urine acid-base parameters were found between SCTR WT and KO (Figure 1A-D) with control water. After 24 hours of base-loading, all three groups responded with significantly increased urinary pH (Figure 1A). Correspondingly, all groups presented with decreased urinary NH_4_^+^ and TA excretion (Figure 1B-D). Interestingly, both WT groups had an initial higher increase in urine pH compared with KOs (Figure 1A).

**Figure 1:**
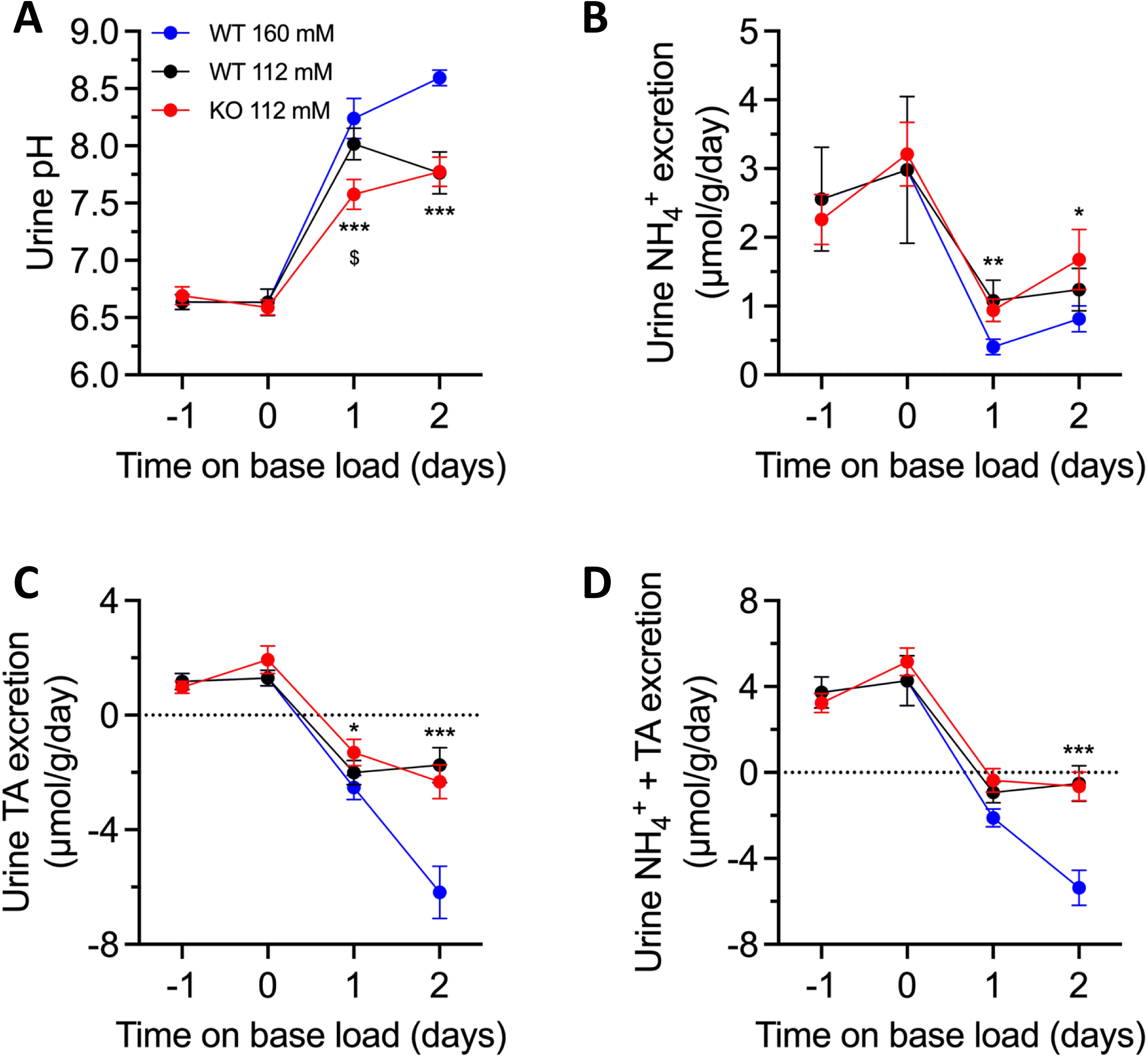
SCTR KO mice respond with diminished urine alkalization upon prolonged base-loading. Urine (A) pH, (B) NH_4_^+^ excretion, (C) titratable acid (TA) excretion and (D) NH_4_^+^ + TA excretion in metabolic cages with control drinking water and after 24 and 48 hours of base-loading with NaHCO_3_-enriched drinking water in SCTR WT mice (112 mM: n=9, black and 160 mM: n=11, blue) and SCTR KO mice (112 mM: n=19, red). Statistical differences were assessed by two-way ANOVA followed by multiple comparisons. NH_4_^+^ excretion was log-transformed before statistical analysis. $: p<0.05 for SCTR KO vs. 112 mM NaHCO_3_ WTs. *, **, and ***: p<0.05, <0.01 and <0.001 for SCTR KO vs. 160 mM NaHCO_3_ WTs, respectively. Error-bars indicate SEM.

Following 48 hours of loading, urine pH increased further in 160 mM NaHCO_3_-loaded WT mice, while urine pH of 112 mM NaHCO_3_-loaded WTs and KOs remained relatively stable (Figure 1A). Concordantly, NH_4_^+^ and TA excretion did not further decrease in 112 mM NaHCO_3_-loaded mice, while TA excretion diminished further in 160 mM NaHCO_3_-loaded WT mice (Figure 1B-D).

Of note, median 24-hour urine output during base-loading was 46% higher in KOs vs WTs receiving 112 mM NaHCO_3_ drinking water, but 7% higher in WTs receiving 160 mM NaHCO_3_ drinking water (Figure S3).

These results indicate that the SCTR plays a role in the regulation of renal acid-base handling during prolonged base-loading. While urine pH only differed initially between WTs and KOs receiving 112 mM NaHCO_3,_ WTs receiving 160 mM NaHCO_3_ (the dose chosen to approximate the higher habitual water intake previously reported in KOs) showed a larger urinary alkalization and a greater diminishment of acid excretion then SCTR KO mice.

### A prolonged base-loading aggravates HCO_3_^-^ accumulation in SCTR KO mice

After examining the urinary response in SCTR WTs and KOs, it was of natural interest to investigate how prolonged base-loading affected systemic acid-base parameters. Venous blood samples were taken from SCTR KO and WT mice at baseline and after 1, 4, and 8 days of base-loading (112 mM NaHCO_3_ for the KOs and 160 mM NaHCO_3_ for the WTs, again in an attempt to adjust for the increased water intake in KOs). WT mice receiving 112 mM NaHCO_3_ were also included for the first 24 hours.

After 24 hours, all three groups presented with increased venous HCO_3_ ^-^ (Figure 2A). 112 mM-loaded KOs had the largest increase, followed by 160 mM-loaded WTs and lastly 112 mM loaded-WTs (Figure 2B). Looking at the systemic response in the 112 mM KO and 160 mM WT groups after 4 and 8 days of base-loading, KOs had higher venous HCO_3_ ^-^ than WTs after 4 days, while no difference was found after 8 days (Figure 3A). Venous blood pH was increased in both groups after 24 hours, but while the WTs gradually normalized their pH towards day 8, venous pH of SCTR KO mice remained elevated (Figure 3B). Venous Cl^-^ decreased in both groups after 24 hours, albeit more in KOs, corresponding to the rise in HCO_3_ ^-^ (Figure 3C). Lastly, both groups presented with an increased pCO which could be interpreted as respiratory compensation, as previously reported to occur in SCTR KO mice during acute HCO_3_^-^ loading (Figure 3D).^14^ This study was not powered for sex-specific analyses, however, the difference between genotypes appeared maintained and, possibly, more pronounced in male mice from day 4 and onwards (Figure S4).

**Figure 2:**
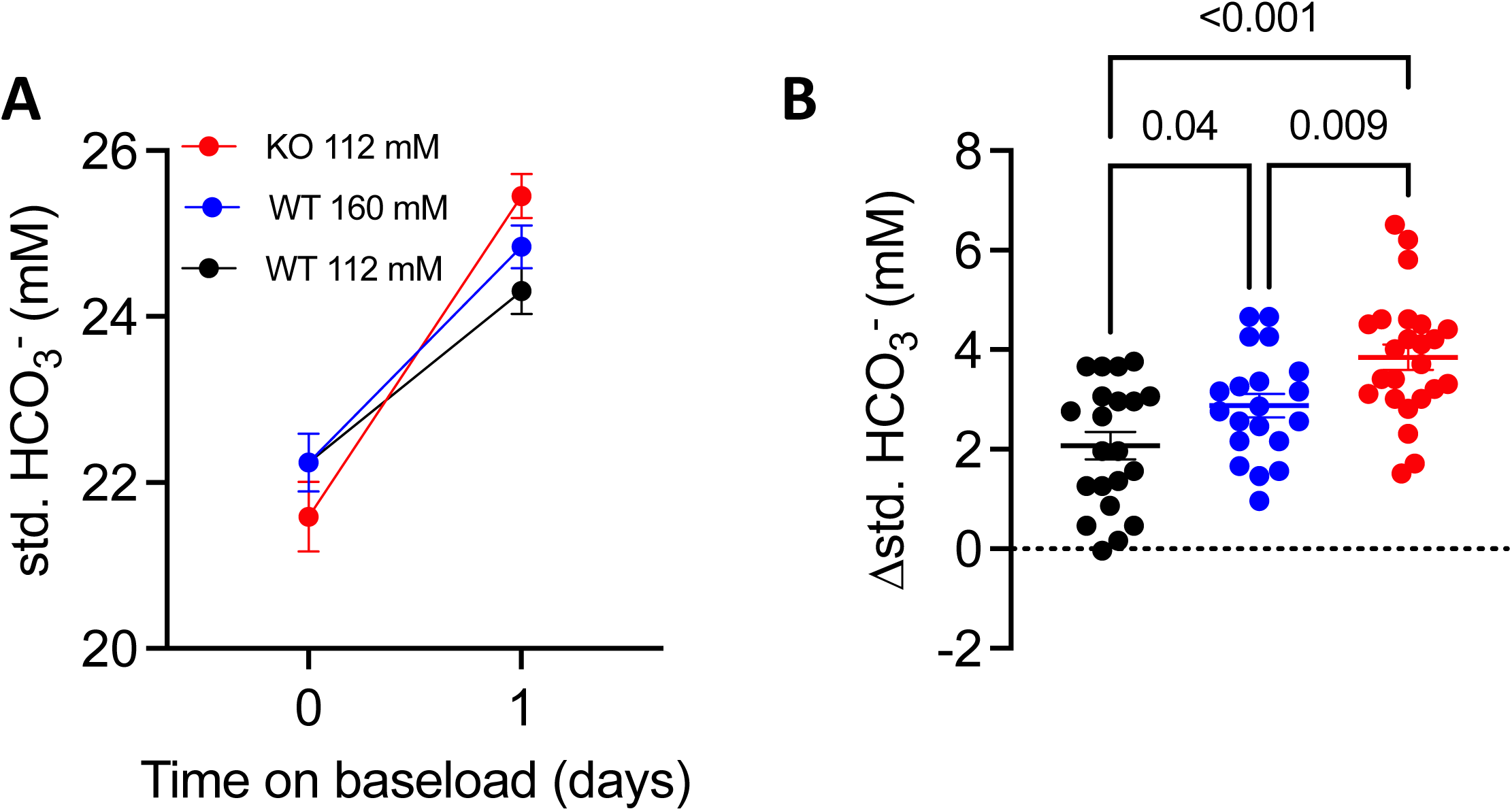
24 hours of base-loading increases venous HCO_3_ ^-^ more in SCTR KO mice compared to SCTR WT mice (A) Venous standard (std.) HCO_3_ ^-^ at baseline and after 24 hours of base-loading with NaHCO_3_-enriched drinking water and (B) the change in venous HCO_3_ ^-^ after 24 hours of base-loading in SCTR WT mice (112 mM: n=21, black and 160 mM: n=15, blue) and SCTR KO mice (112 mM: n=23, red). Statistical differences were assessed by one-way ANOVA followed by multiple comparisons (B). Error-bars indicate SEM.

**Figure 3:**
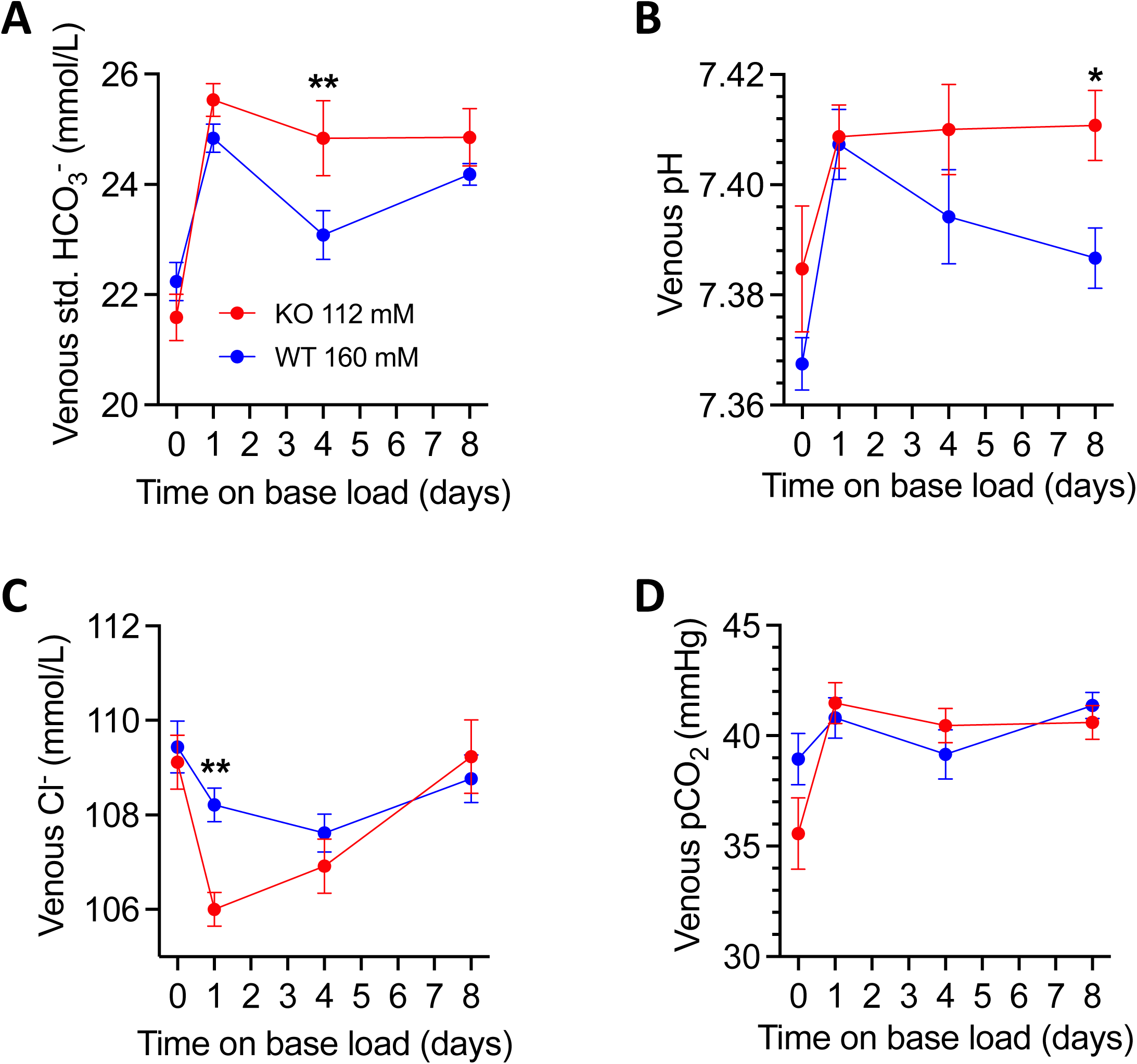
SCTR KO mice exhibit increased base retention during prolonged base-loading Venous (A) Std. HCO_3_^-^, (B) pH, (C) Cl^-^ and (D) pCO_2_ at baseline and after 1, 4 and 8 days of base-loading with NaHCO_3_-enriched drinking water in SCTR WT mice (160 mM: n=15-17, blue) and SCTR KO mice (112 mM, n=11-23, red). Statistical differences between the two groups were assessed by two-way ANOVA followed by multiple comparisons. Error-bars indicate SEM.

To further expand on the renal adaptation to base-loading, urinary acid excretion (NH_4_^+^ + TA) as a function of venous standard HCO_3_^-^ was assessed at baseline and after 24 hours of oral base-loading (Figure 4). Both WT groups (112 mM: black, 160 mM: blue) exhibited a sharp decline in urinary acid excretion after base-loading, eventually reaching a positive net base excretion and a plasma HCO_3_^-^ of 24.3 mmol/L. This relationship was blunted in the KO mice (slope significantly less steep, p=0.003) who reached a lower net urinary base excretion despite having a higher venous HCO_3_^-^ compared to WTs. Similar trends were observed in sex-stratified analyses (p=0.012 in females, p=0.11 in males).

**Figure 4:**
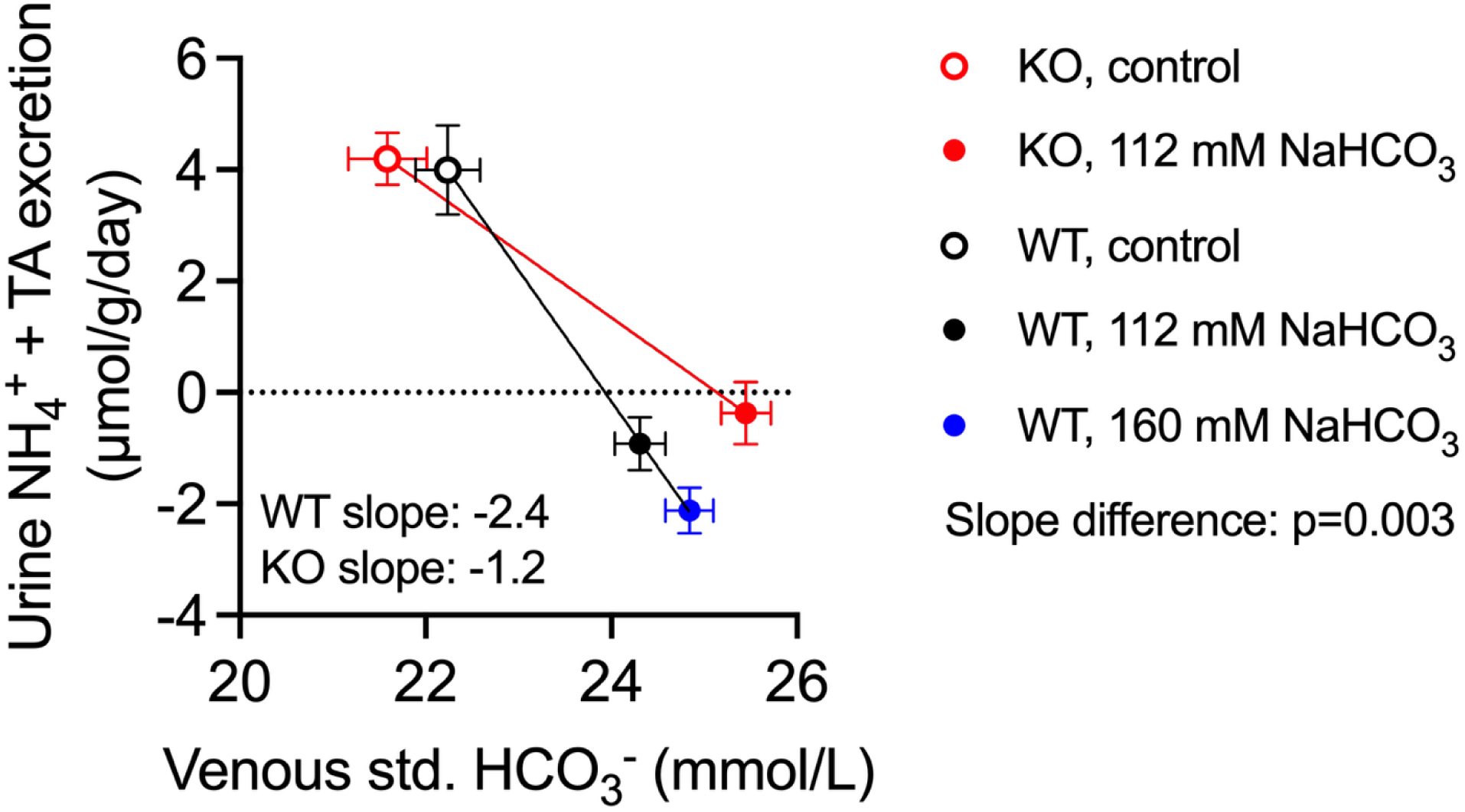
SCTR KO mice reach a negative acid excretion at higher venous HCO_3_^-^ concentrations Urinary acid (NH_4_^+^ + titratable acid) excretion as a function of venous standard (std.) HCO_3_ ^-^ in SCTR WT and SCTR KO mice. Data are combined from figure 1 and figure 2. Note that a negative total NH_4_^+^ and TA excretion equates a positive net excretion of base. Error-bars indicate SEM.

### SCTR KO mice fail to adequately increase pendrin function despite aggravated alkalosis

We recently found that in isolated perfused cortical collecting ducts, secretin pre-incubation (10 nmol/L for 10 min) increased pendrin function ∼2-fold in WTs but had no effect in SCTR KO tubules.^14^ With this in mind, we investigated pendrin protein abundance and function in SCTR KO and WT mice during control conditions and following 24 hours of base-loading with 112 mM NaHCO_3_-enriched drinking water. SCTR KO and WT mice displayed similar pendrin abundance under control conditions (Figure 5A-B and Figure S5) and following 24 hours of base-loading, pendrin abundance increased equally in SCTR KO and WT mice (Figure 5A-B and Figure S5). These results show that SCTR KO mice were able to fully increase pendrin protein abundance after a 24-hour base-challenge.

**Figure 5:**
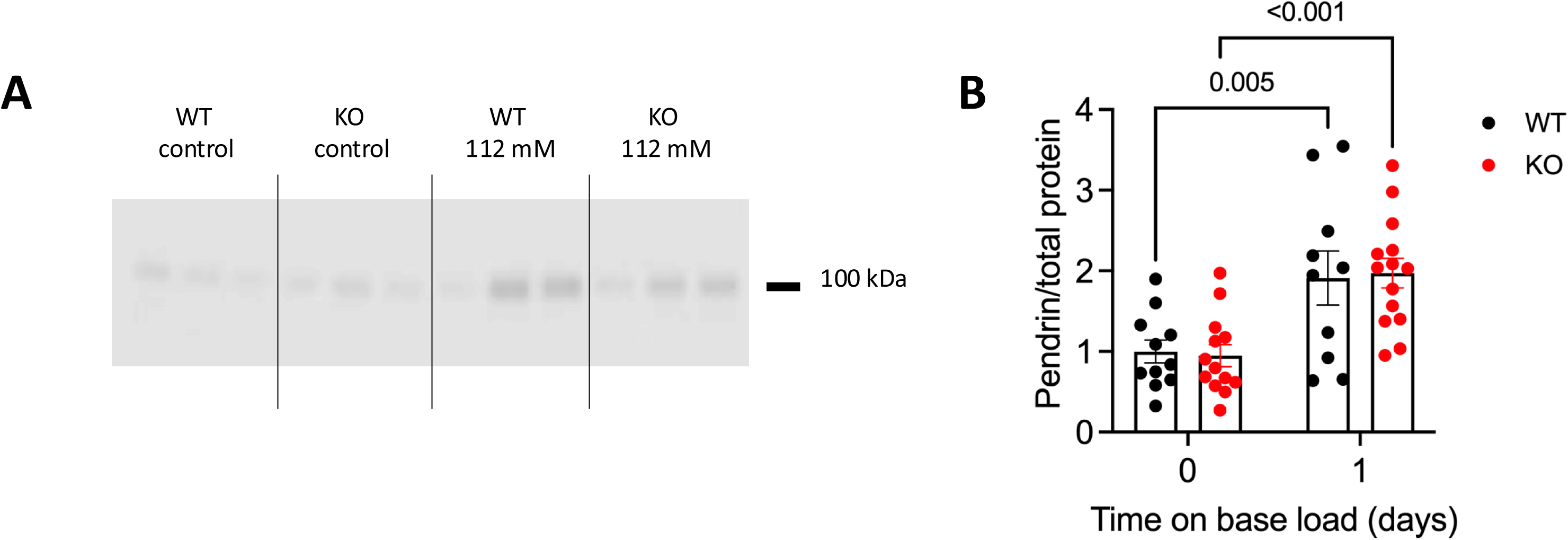
SCTR KO mice show normal pendrin protein abundance regulation during prolonged base-loading. (A) original blot showing protein abundance of pendrin in WT and KO mice under resting conditions and after 24 hours of base-loading with NaHCO_3_-enriched drinking water (112 mM). Uncropped full-length blots are shown in Figure S5. (B) quantification of pendrin protein abundance in the SCTR WT (n=10-11) and KO mice (n=13-14) during resting conditions and after 24 hours of base-loading with NaHCO_3_-enriched drinking water (112 mM). Pendrin protein abundance was normalized to total loaded protein (using Coomassie staining)^34^ and expressed relative to SCTR WT mice during control conditions. Statistical differences were assessed by two-way ANOVA followed by multiple comparisons (panel B). Error-bars indicate SEM.

Subsequently, using tubule perfusion and individual beta-IC intracellular pH imaging, we investigated pendrin function in isolated collecting ducts from SCTR WT and KO mice under control conditions and after 24 hours of base-loading with 112 mM NaHCO_3_-enriched drinking water. WTs as compared to KO beta-ICs exhibited a ∼2-fold higher increase in the intracellular alkalization rate when applying the luminal Cl^-^ removal maneuver (Figure 6A-C). After base-loading resting pH_i_ in beta-IC was lower in both WT and KO beta-ICs. No resting pH_i_ differences were observed between WT and KO beta-ICs during control conditions or after 1 day of base-loading. To take into account that baseline pH_i_ could affect the initial alkalization rate upon luminal chloride removal, we analyzed the intracellular alkalization rate as a function of baseline pH_i_, which confirmed the higher alkalization rate in WT with a significantly different y-axis intercept of the two lines (using a common slope coefficient, Figure 6E). Derived from previously measured buffer capacity in single beta-ICs,^32^ we could estimate the cellular buffer capacity β_i_, the β_HCO3_- and hence the total buffer capacity, β_total_. This permitted the calculation of the apparent HCO_3_ ^-^ flux over a range of different pH_i_ values in WT and KO pendrin positive ICs. The results showed increases in pendrin-positive IC base flux in WT and KO cells, but with a markedly lower base flux in SCTR KO cells after 24h base loading (Figure S2).

**Figure 6:**
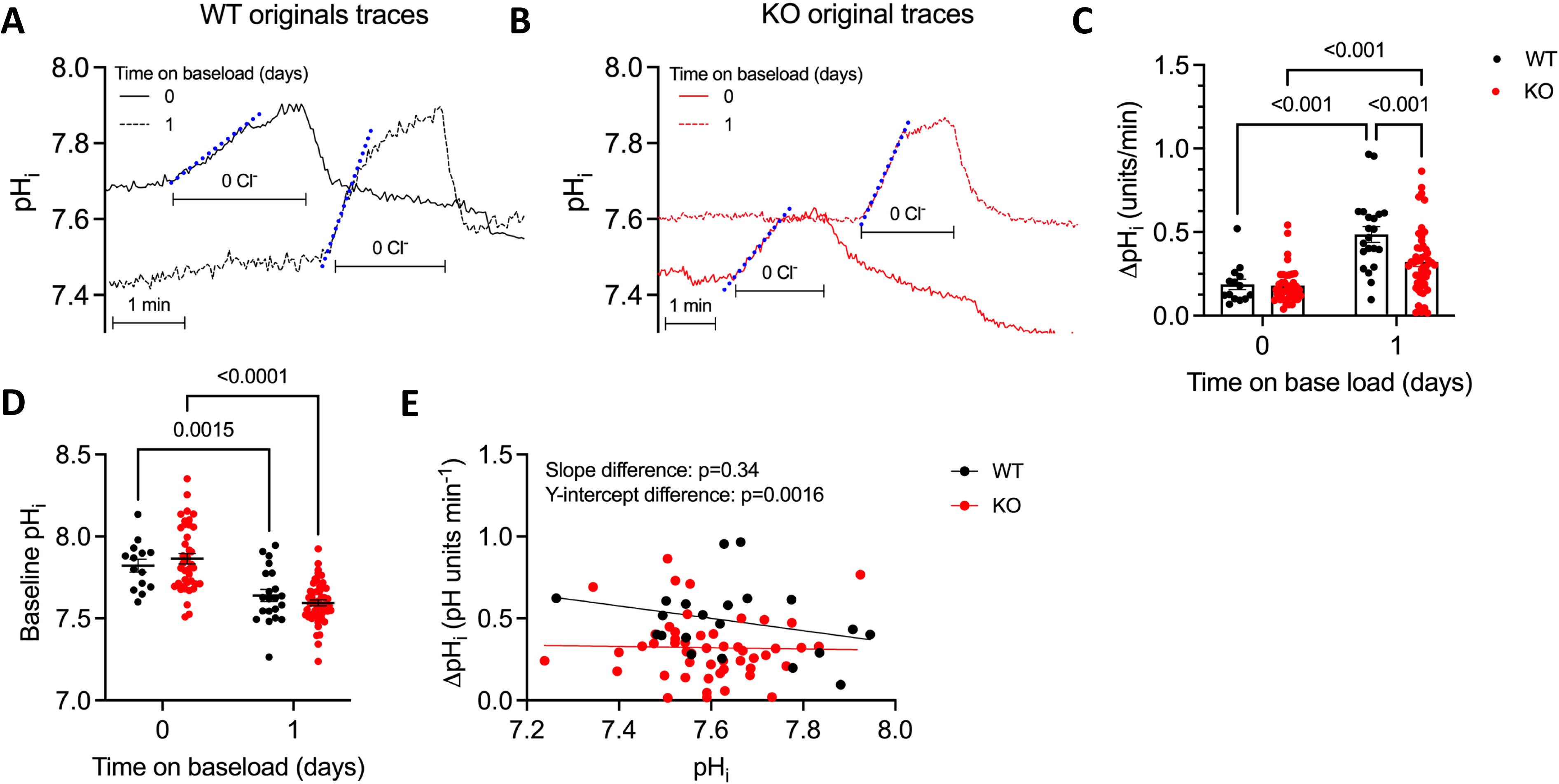
SCTR KO mice exhibit impaired pendrin function regulation during prolonged base-loading (A+B) original traces depicting intracellular pH as a function of time in isolated perfused cortical collecting ducts of SCTR WT mice (A) and SCTR KO mice (B). (C) Summary of pendrin function in isolated perfused cortical collecting ducts from SCTR WT and KO mice during control conditions and after 24 hours of base-loading with NaHCO_3_-enriched drinking water (112 mM). (D) Baseline (i.e. before luminal Cl^-^ removal) intracellular pH in SCTR WT and KO pendrin positive cells in control and base-loaded animals. (E) the association between baseline intracellular pH and the acute intracellular alkalization rate upon luminal chloride removal in base-loaded SCTR WT and KO mice. Pendrin function was assessed as the initial rate change of intracellular pH after luminal chloride removal (graphically illustrated by the blue line in panels A+B). Statistical differences between pendrin function and baseline intracellular pH assessed by two-way ANOVA followed by multiple comparisons (panel C+D), slope differences (panel E) was assessed by testing if there was a significant interaction between genotype and intracellular pH on the association between intracellular pH and pendrin function (F-test, panel E), genotype differences were also addressed by assessing the effect of genotype on pendrin function after adjusting for baseline intracellular pH (F-test, panel E). Error-bars indicate SEM.

These results indicate that the capacity to increase pendrin function during chronic base-loading (assessed after 24 hours of 112 mM NaHCO_3_ loading) depends significantly on the presence of functional SCTR. The fact that SCTR KO mice were able to fully increase pendrin protein abundance and partially increase pendrin function illustrates that other mechanisms besides secretin influence the regulation of pendrin expression and function.

### A higher arterial HCO3 - is associated with higher plasma secretin and renal SCTR levels after 24 hours of base or acid-loading

The role of secretin and its receptor in the systemic regulation of plasma HCO_3_ ^-^ levels following an acute metabolic alkalosis was previously examined by our group. Here, an acute oral NaHCO_3_-gavage significantly increased plasma secretin levels.^13,37^

We therefore investigated whether plasma secretin levels and renal SCTR expression were modulated by systemic acid-base status during prolonged acid/base-loading. As plasma secretin has a very short half-life and anesthesia via ketamine/xylazine suppresses respiration and thus induces respiratory acidosis, normal ventilation was maintained by mechanical ventilation during sampling of plasma and kidney tissue from acid- and base-loaded mice. After 24 hours of acid/base-loading, plasma secretin levels were higher in base-loaded compared with acid-loaded animals (Figure 7A). Assessment of the relationship between arterial HCO_3_^-^ and plasma secretin levels revealed a borderline significant association (p=0.06, Figure 7B), which was stronger in females (p=0.009, Figure S6). When examining the relative SCTR expression in kidney tissue from the same mice, a similar relationship appeared, i.e. SCTR expression was significantly higher in base-loaded than acid-loaded animals (Figure 7C and Figure S7). Here, a higher arterial HCO_3_^-^ was associated with a significantly higher renal SCTR expression (Figure 7D).

**Figure 7:**
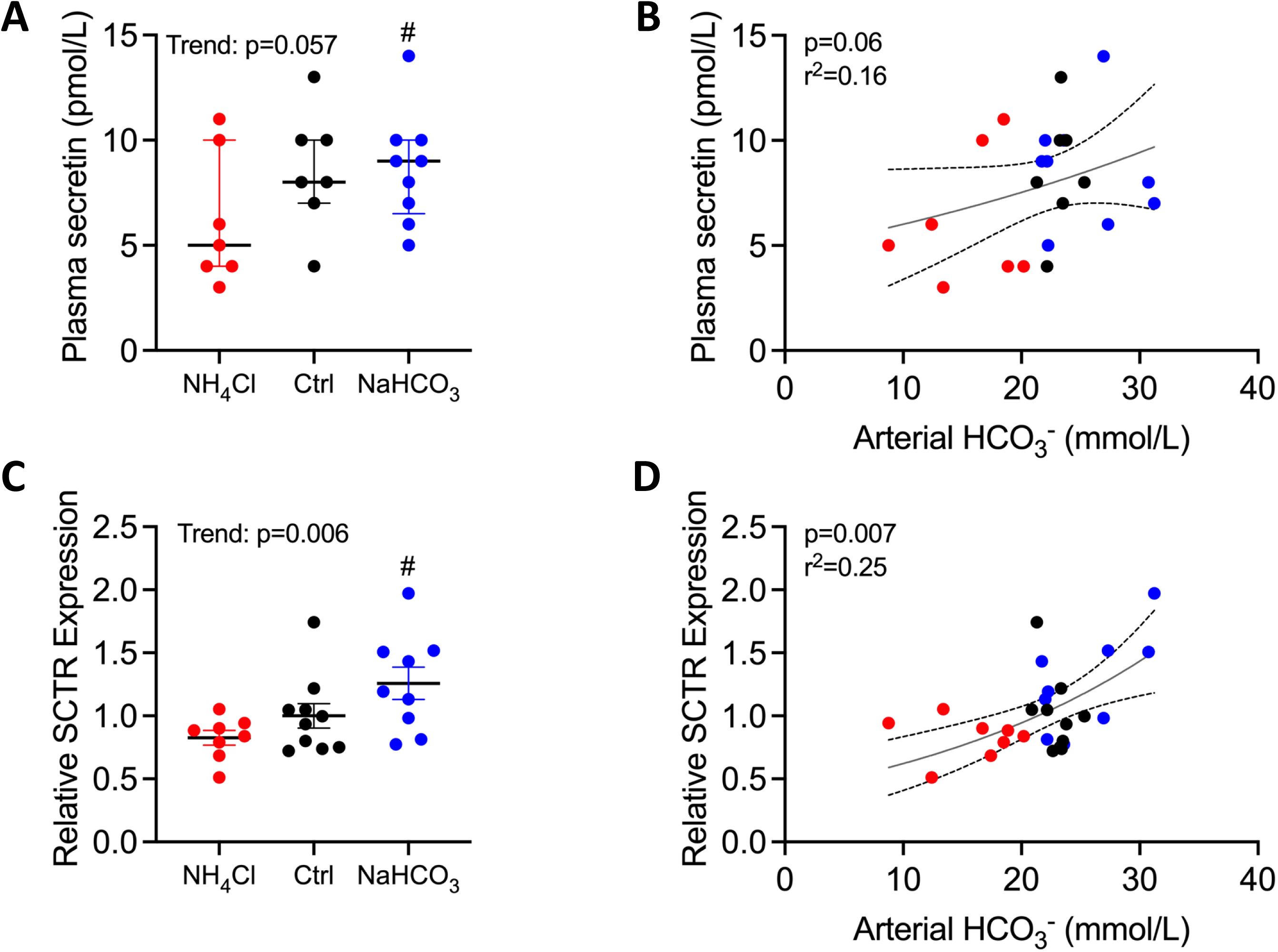
Plasma secretin and renal SCTR expression levels are modulated by acid/base-loading. (A) Plasma secretin (pmol/L) in C57BL/6 mice under control conditions (n=7, black dots), after 24 hours of acid-loading (200 mM NH_4_Cl, n=7; red dots) and after 24 hours of base-loading (200 mM NaHCO_3_, n=9, blue dots). (B) Plasma secretin (pmol/L) as a function of arterial HCO_3_^-^ (mmol/L) across the same three groups (n=23). (C) SCTR mRNA expression in kidney tissue from the same three groups (control: n=10, 200 mM NH_4_Cl: n=8, 200 mM NaHCO_3_: n=9). SCTR mRNA expression was normalized to the mean of the control group. (D) Relative SCTR expression as a function of arterial HCO_3_^-^ (mmol/L) across the same three groups (n=27). SCTR expression was normalized to that of two housekeeping genes (mPPIA and mHPRT) and expressed relative to that of control mice. Statistical differences were assessed using one-way ANOVA followed by multiple comparisons (panel A+C) or linear regression (panel B+D). In one-way ANOVA analyses, a test for trend, i.e. linear trends in plasma secretin/kidney SCTR levels from NH_4_Cl-loaded to control-loaded to NaHCO_3_-loaded animals. Plasma secretin and kidney SCTR levels were log10-transformed before statistical analysis. #: p<0.05 vs. acid-loaded animals. Data are presented as medians with IQR (panel A) or mean with SEM (panel C). Error-bars indicate 95% CI of the regression line in panel B+D.

These results indicate a functional link between secretin, its receptor and systemic acid-base balance where systemic acid-base status/the need for base excretion modulates plasma secretin levels and renal SCTR expression.

## Discussion

We previously described that secretin plays a significant role in the acute renal compensation of a metabolic alkalosis.^14^ This raised the question whether secretin and its exerted effect on the SCTR are also important during prolonged base-loading and how changing needs for renal acid/base excretion affect plasma secretin levels and kidney SCTR expression.

We found that base-loading sharply increased urinary pH in both SCTR KO and WT mice with a corresponding decrease in NH_4_ ^+^ and TA excretion. However, the ability to alkalize the urine was reduced in KO compared with WT mice.

When assessing systemic acid-base balance during base-loading, both WT and KO mice had a significant increase in blood HCO_3_ ^-^ levels (3-4 mmol/L) after 1 day of base-loading. However, whereas WT mice were able to almost fully correct the alkalosis, returning HCO_3_^-^ almost to baseline levels after 4 days, KO mice exhibited no clear correction of the base excess. Consistent with this, KO mice reached higher blood HCO_3_^-^ levels before achieving a positive net base excretion. This indicates that KO mice reach steady-state conditions at significantly higher blood HCO_3_^-^ levels than WTs. However, after 8 days of base-loading, no clear differences in venous HCO_3_^-^ concentrations were found between SCTR WT and KO mice. This could indicate that the importance of the SCTR for acid-base handling is most pronounced during the initial days of increased base intake. In sex-stratified analyses, the difference between genotypes appeared maintained and possibly more pronounced in male mice. Noteworthy, a recent study found that male mice have larger increases in blood bicarbonate levels when base-loaded as compared to female mice.^38^ As such, male mice, that appear more susceptible to developing metabolic alkalosis, may exhibit a prominent phenotype when components of renal acid-base handling are defunct. The increase of venous HCO_3_ ^-^ in WT mice from day 4 to day 8 could be a consequence of the hypertonic drinking water or higher NaHCO_3_ intake in WTs at the high dose, as addressed later in this discussion.

When examining the protein abundance and function of pendrin, it became clear that the SCTR plays a pivotal role in the upregulation of pendrin function (measured after 24 hours of 112 mM NaHCO_3_ loading) but that other mechanisms partake in the regulation of protein abundance. Here, it is worth mentioning that the basolateral Na^+^-dependent Cl^-^/HCO_3_ ^-^ exchanger AE4 (SLC4A9) is pivotal for the ability to regulate pendrin protein expression during both alkalosis and acidosis.^32^ The underlying mechanism for SLC4A9-dependent regulation of pendrin remains elusive. However, WNK-signaling has been shown to affect pendrin regulation, and missense mutations in WNK4 (that result in increased WNK4 expression) cause overactivation of pendrin^39^. In the distal convoluted tubule, low intracellular chloride levels lead to activation of the WNK4-SPAK pathway which increases NCC transport activity via phosphorylation.^40^ As AE4 is thought to be a Cl^-^ exit pathway, it is possible that AE4 could participate in pendrin regulation by modulating intracellular Cl^-^ concentrations. Whether secretin signaling affects AE4 abundance or function remains to be determined. Our data indicates that the mechanisms governing pendrin protein abundance during base-loading appear unaffected by loss of secretin signaling, but that without secretin signaling the ability to fully activate pendrin is hampered.

Finally, we found that after 24 hours of acid/base-loading, both plasma secretin levels and renal SCTR expression were higher in base-loaded compared to acid-loaded animals. It should be noted that no significant differences were found between control and base-loaded animals. Even so, a higher arterial HCO_3_ ^-^ concentration associated with higher renal SCTR expression levels (p=0.007) and borderline significantly associated with higher plasma secretin levels (p=0.06). Our results indicate a graded renal SCTR response and, possibly, a graded plasma secretin response across a wide range of arterial HCO_3_^-^ values, including the acidotic range. A decrease in secretin-mediated HCO_3_^-^ secretion would be meaningful during conditions of acid excess, as constitutively activated beta-IC HCO_3_^-^ secretion aggravates metabolic acidosis.^32,39^ Supporting this, among the NH_4_Cl-loaded C57Bl/6J mice, the mice with the highest pendrin expression after acid-loading exhibited the most severe acidosis (Figure S8-S9). Whether decreases of plasma secretin/renal SCTR expression are physiologically relevant during acid excess is an intriguing research question but not addressed in this study.

Our results underline that secretin and its exerted effects extend beyond post-prandial gastrointestinal regulation, and that the hormone also plays a role in the renal regulation of acid-base homeostasis. An important question is the origin of the released secretin and the exact triggering mechanisms. We have previously found that secretin release from the small intestine is increased during a mimicked metabolic alkalosis,^14^ however, in those experiments, no reduction of plasma secretin release was observed during acidotic conditions. One could speculate that acidosis reduces the half-life of plasma secretin, possibly mediated through changes in plasma protease activity, thus decreasing circulating levels of the hormone.

This study has several limitations. Notably, SCTR KO mice had global deletion of the SCTR, which should be viewed in context of its expression in the lungs and gastrointestinal tract.^16^ It is possible that the secretin response is a feed-forward signal from the intestine to the kidney and that intravenous administration of HCO_3_ ^-^ or an imposed respiratory alkalosis could produce different results. We have previously tried to address this issue via acute intraperitoneal administration of HCO_3_^-^, which produced results similar to those obtained after oral intake.^14^ Whether this translates to a chronic setting remains unknown. However, if lack of SCTR expression in the lungs, intestine or pancreas was the cause of the aggravated base retention in SCTR KOs, and there was no renal phenotype, one would expect a higher base excretion in the KOs. Hence, the concurrently lower base excretion and larger increase in blood base levels in KOs (at 112 mM in KOs vs. 160 mM WTs), supports that the observed phenotype is caused by impaired renal base excretion.

It should also be mentioned that the used NaHCO_3_ doses (112-160 mmol/L) are at the lower end of what other laboratories often use (several studies use 280 mmol/L).^32,41^ We used 112 mmol/L, because we previously have shown that beta-IC dysfunction (KO of pendrin or CFTR) causes base retention even at this moderate degree of base-loading.^19^ Further, a clear advantage is that we avoid providing excessively hypertonic drinking water that could induce a variety of confounding effects (e.g., dehydration). A limitation is that the observed increases in blood HCO_3_ ^-^ levels are quite modest. An alternative approach could have been to introduce base-loading by adding base equivalents to the chow.

The difference in the administered NaHCO_3_ concentration between the KO and WT group also needs mentioning. The 160 mM solution given to the WTs is slightly hypertonic compared with plasma, which could have several consequences. Firstly, intracellular dehydration is a natural result of hypertonic plasma, which could result from consumption of the 160 mM solution. This might activate the renin-angiotensin-aldosterone-system, potentially increasing urinary H^+^-excretion and aggravating the systemic base retention. Furthermore, an increased amount of filtered sodium delivered to the nephron could also possibly drive H^+^-excretion via sodium-proton exchangers (i.e., NHE3). However, both these effects would aggravate the alkalosis in WT mice, diminishing the observed differences between genotypes. Another thing to note is the urine output in the experimental groups, which would be expected to associate with water intake. As expected, we observed higher urine outputs in SCTR KO than WT mice during 112 mM NaHCO_3_-loading. However, WT mice exposed to 160 mM NaHCO_3_ drinking water had a significantly higher urine output than WTs with 112 mM NaHCO_3_, and a similar urine output compared to SCTR KO mice with 112 mM NaHCO_3_ drinking water. The cause of this is uncertain, but could be due to the hypertonicity, which could increase water intake. Assuming that the mice are in steady-state after two days of base-loading, the fact that SCTR WT mice exposed to 160 mM NaHCO_3_ water excreted more base equivalents than SCTR KO mice exposed to 112 mM NaHCO_3_ drinking water is in alignment with a higher NaHCO_3_ ingestion in SCTR WT mice at these doses. It is possible that these effects (intracellular dehydration, RAAS activation and increased base intake) are responsible for the increase in venous HCO_3_ ^-^ in WT mice from day 4 to day 8 of base-loading.

Lastly, we only measured pendrin protein expression and activity after 24 hours of 112 mM NaHCO_3_-loading. It is possible that a higher pendrin protein abundance in WTs would be revealed by increasing the base intake to match the intake in the KOs. However, the mean increase in pendrin protein abundance of WT and KO mice was ∼2-fold, which is equal to or larger than what has previously been found using significantly larger doses.^32,41^ It is worth noting that SCTR KO mice exhibited markedly lower pendrin function despite higher blood HCO_3_ ^-^ and lower blood Cl^-^ levels, both of which are potent stimulators of beta-IC activity and pendrin function.^42,43^ It is also possible that a larger difference in pendrin activity would have been found at day 4, where the largest difference in venous HCO_3_^-^ levels were found. In summary, we found that the secretin/SCTR system plays an important role in the alleviation of prolonged base intake in mice. Additionally, systemic acid-base status appears to modulate kidney SCTR mRNA and, possibly, plasma secretin levels. These findings support our overarching suggestion that secretin is a bona fide HCO_3_^-^ excretory hormone relevant for the maintenance of acid-base homeostasis Data Availability Statement: The data that support the findings of this study are available from the corresponding author upon reasonable request.

## Acknowledgements

We thank Karen Skjødt Sørensen for expert technical assistance.

## Author contributions

Conceptualization: PB, HV

Methodology: SLS, JL, MVS, PB, LWT, JJH, IMM, BH, HV

Data curation: TJ, JFA, LWT, MVS, PB, IMM, BH

Formal analysis: TJ, PB

Supervision: SLS, JL, PB

Funding acquisition: JL

Visualization: TJ, JFA, PB

Project administration: TJ, PB

Writing – original draft: TJ, JFA, PB

Writing – review & editing: TJ, JF, SLS, MVS, JL, PB, IMM, JJH, BH, HV

