## Supplemental Material for "Impaired renal base excretion in secretin receptor knock-out mice during prolonged base-loading"

### Supplementary Figures

#### Figure S1

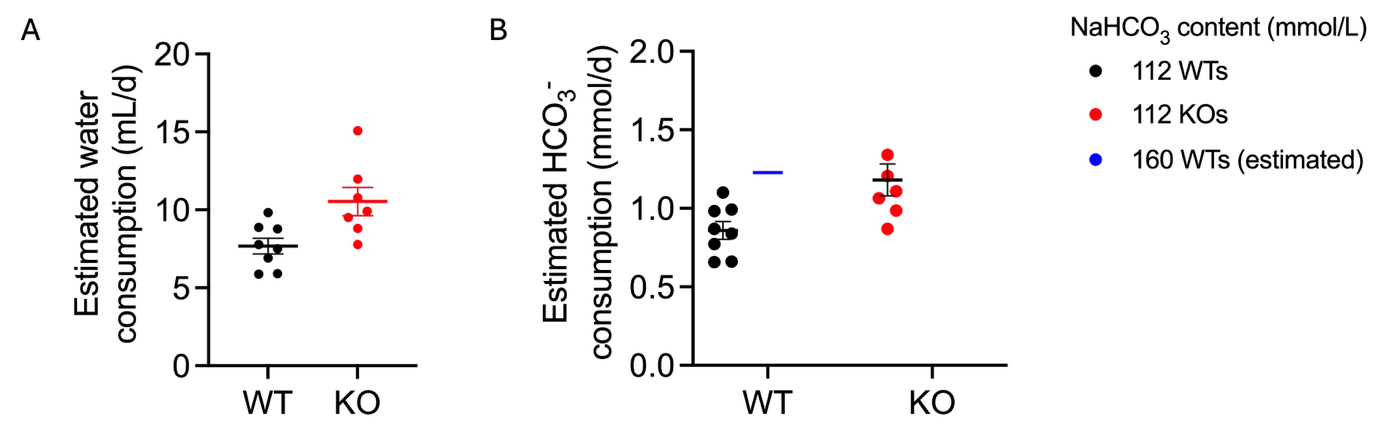

A) Estimated water intake in SCTR WT and KO mice in metabolic cages with 112 mM NaHCO_3_ drinking water. Water intake was estimated by the daily change in water bottle weight. B) estimated NaHCO_3_ consumption in SCTR WT and KO mice during exposure to 112 mM NaHCO_3_ (WTs and KOs) and 160 mM NaHCO_3_ drinking water.

#### Figure S2

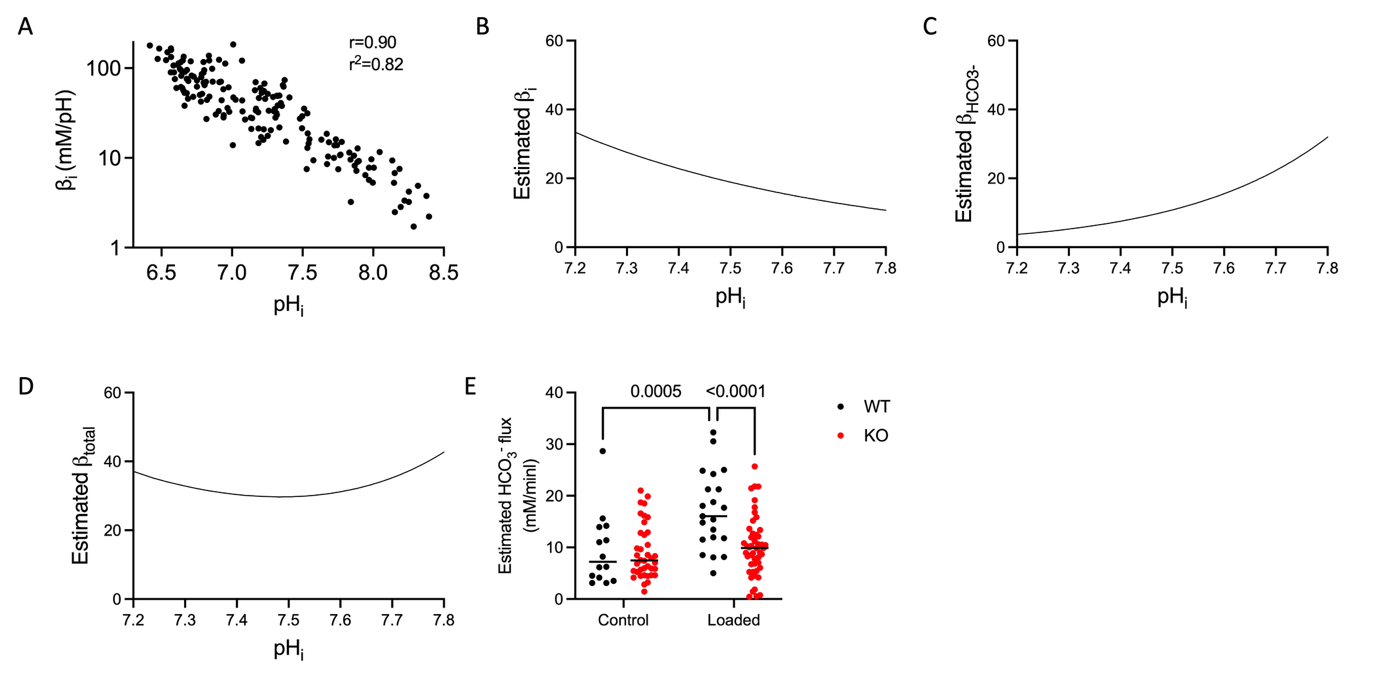

A) Association between intracellular pH_i_ and intrinsic buffer capacity (replotted from Vitzthum et al^1^) showing a very strong association between log10 transformed intrinsic buffer capacity and intracellular pH. B) Estimated intrinsic buffer capacity in the range of intracellular pH of 7.2 to 7.8 using data from panel A. C) estimated contribution of intracellular bicarbonate to total buffer capacity in the range of intracellular pH of 7.2 to 7.8. D) Estimated total buffer capacity as a function of intracellular pH using panel B+C. E) estimated base flux per cell using dpH_i_/min and estimated total buffer capacity in STCR WT and KO mice during control conditions and after 24 hours of exposure to 112 mM NaHCO_3_ drinking water. Statistical differences were assessed by two-way ANOVA followed by multiple comparisons.

#### Figure S3

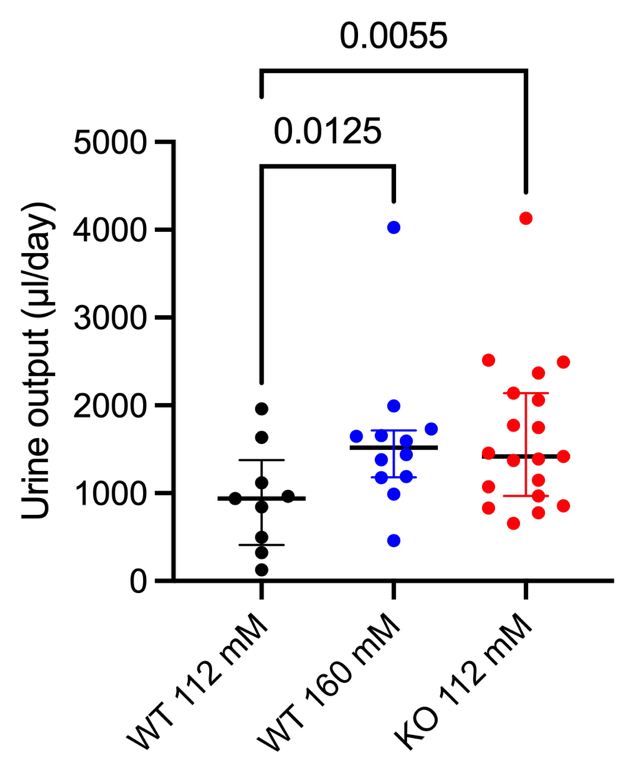

Mean volume of collection urine during two days of base-loading with 112 mM NaHCO_3_ drinking water in SCTR WTs (black) and KOs (red) and 160 mM NaHCO_3_ drinking water in SCTR WTs (blue). Statistical differences were assessed by one-way ANOVA followed by multiple comparisons.

#### Figure S4

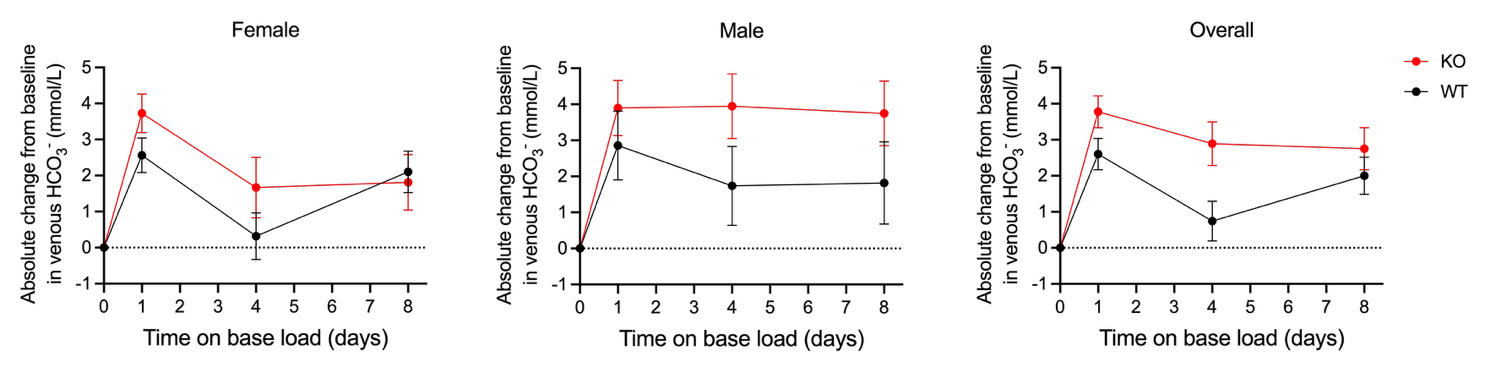

Sex stratified (left and middle panels), and sex adjusted analyses (right panel) of changes in venous HCO_3_^-^ during prolonged NaHCO_3_-loading with NaHCO_3_-enriched drinking water (160 mM for WTs, 112 mM for KOs). Estimates were derived from a linear mixed-effects model for repeated measures with venous HCO_3_^-^ as outcome and sex, time, genotype, sex-by-time interaction, genotype-by-time interaction, and sex-by-time-by-genotype interaction as fixed effects and mouse ID as random intercept and time as a random slope.

#### Figure S5

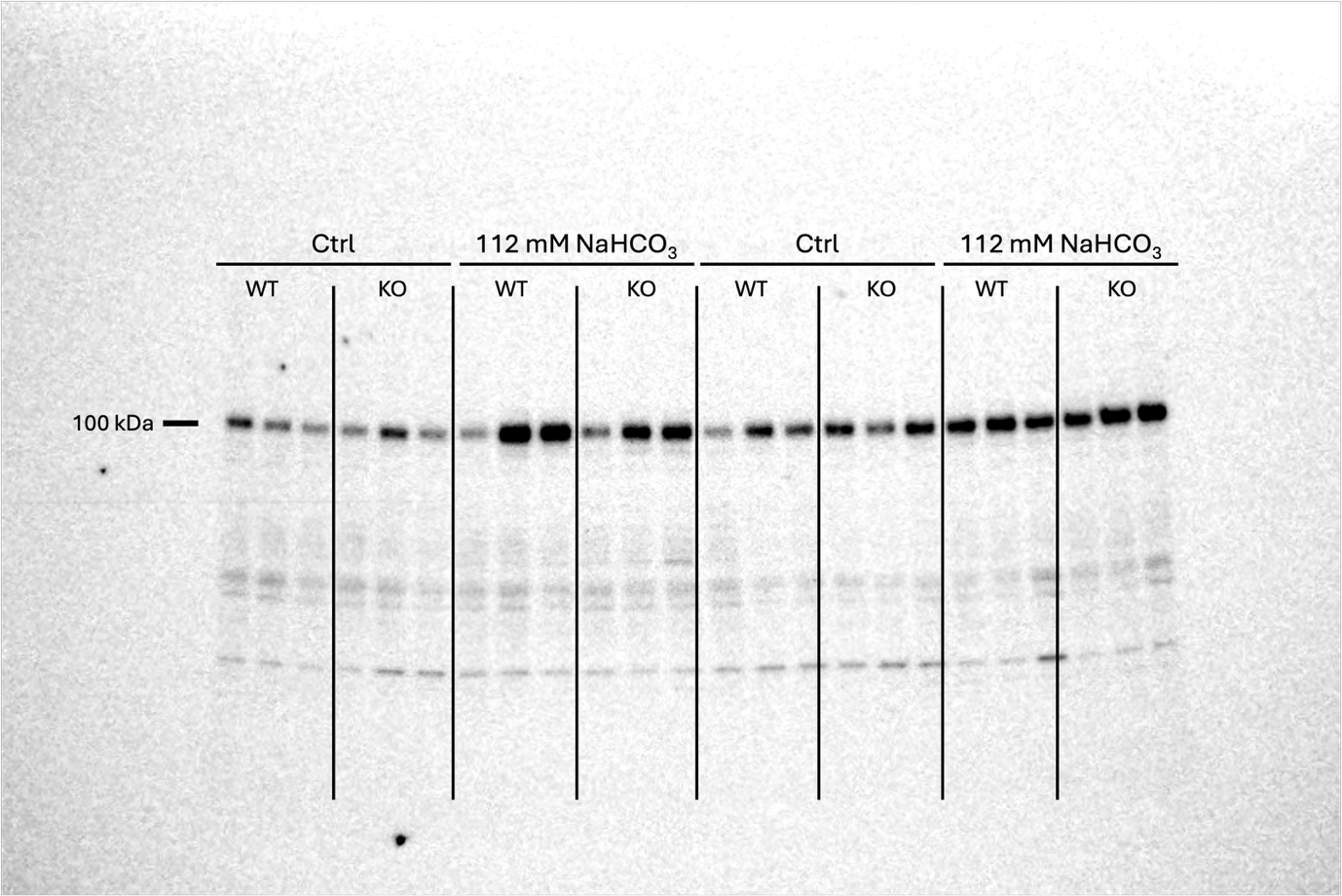

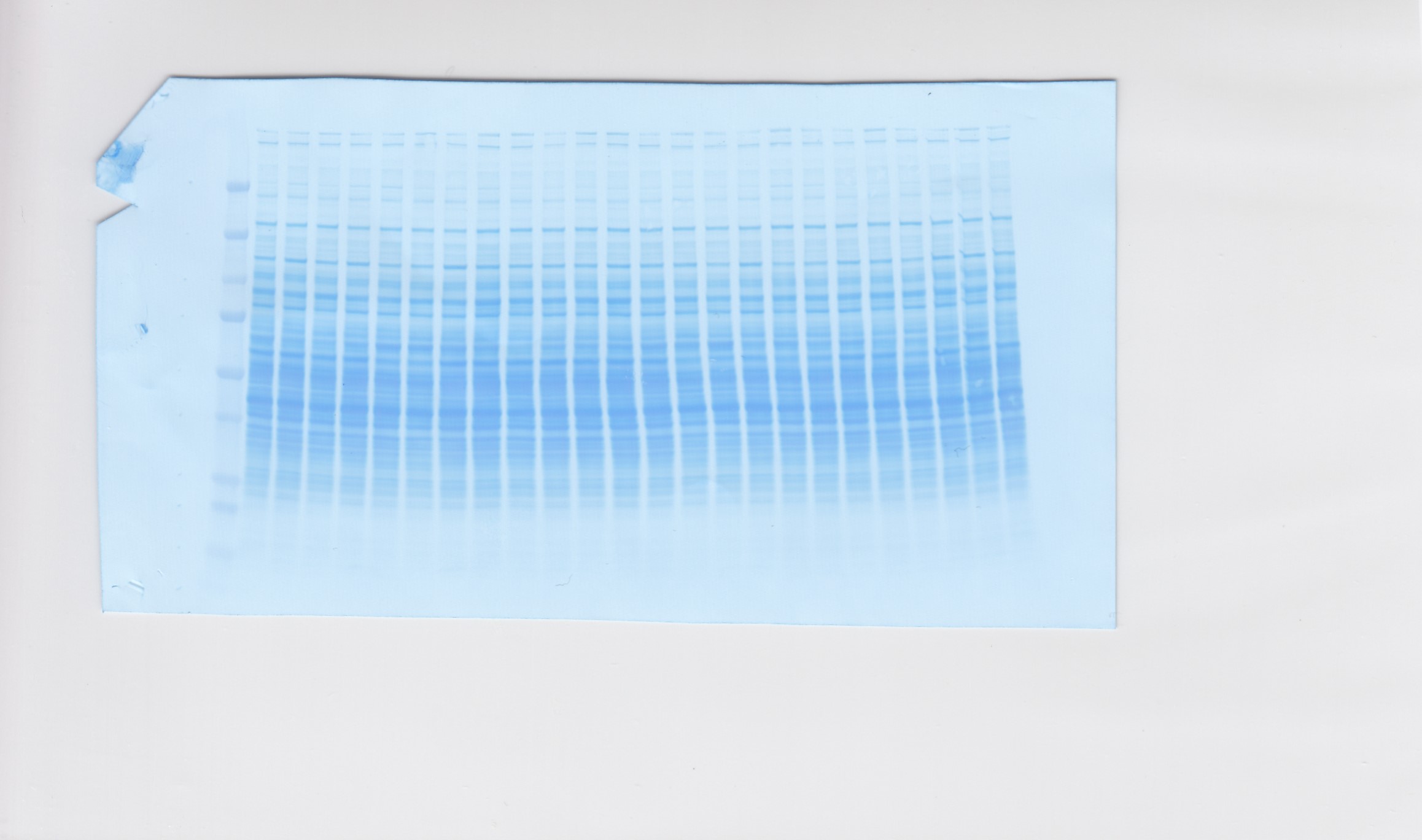

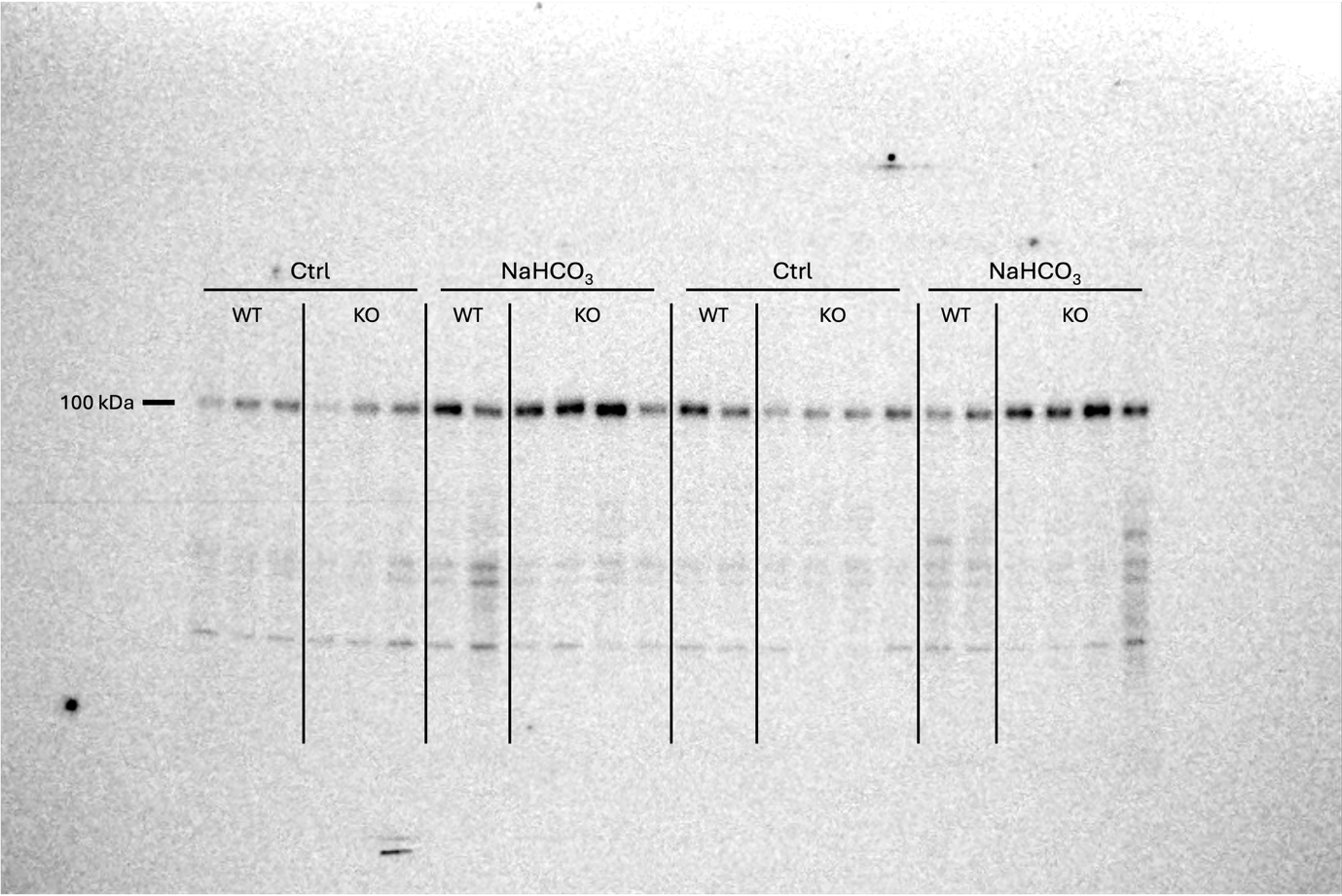

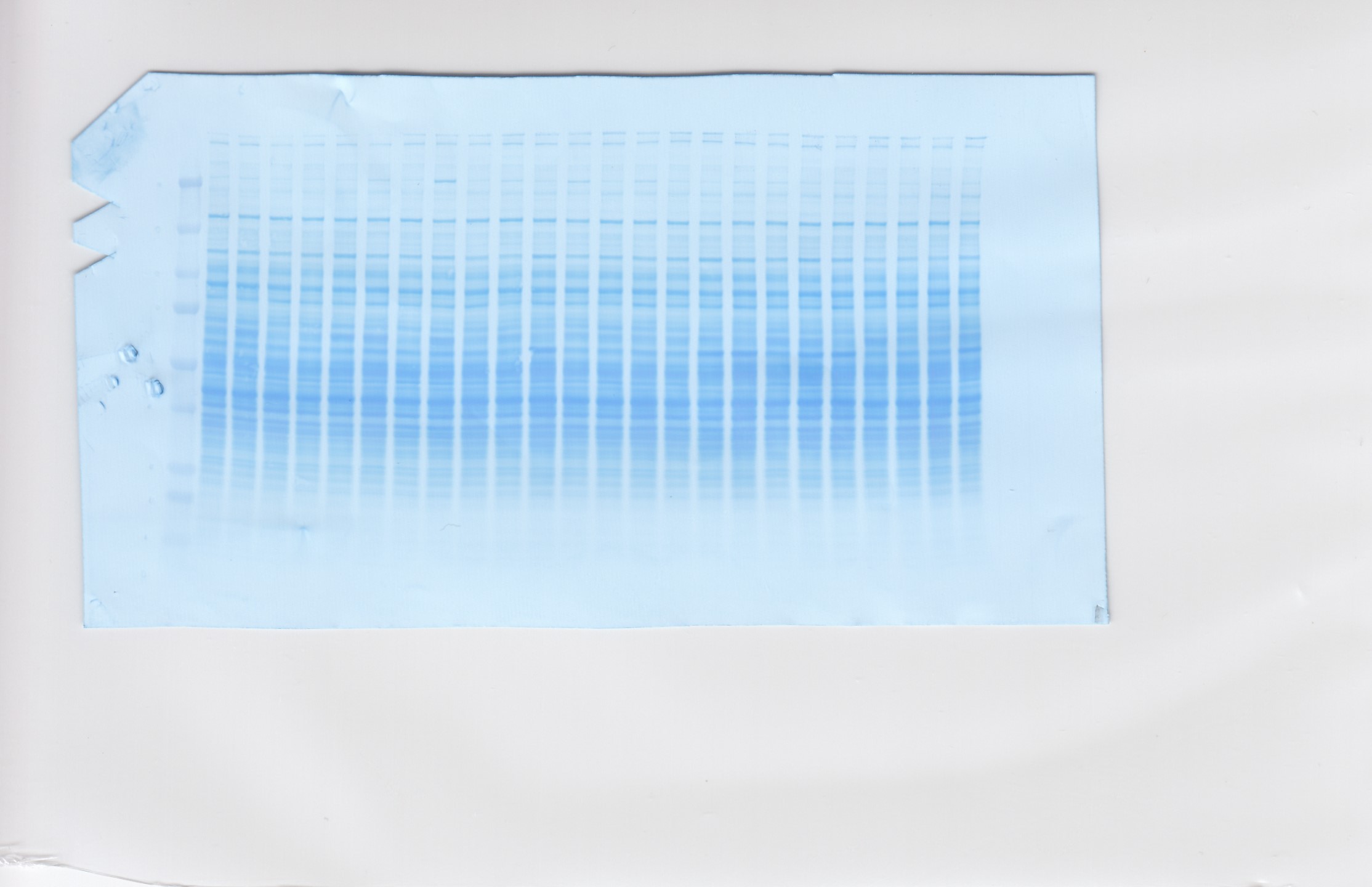

Uncropped full-length pendrin blots and corresponding Coomassie stainings of the membranes of kidney lysates from SCTR WT and KO mice exposed to 24 hours of either control or 112 mM NaHCO_3_ drinking water.

#### Figure S6

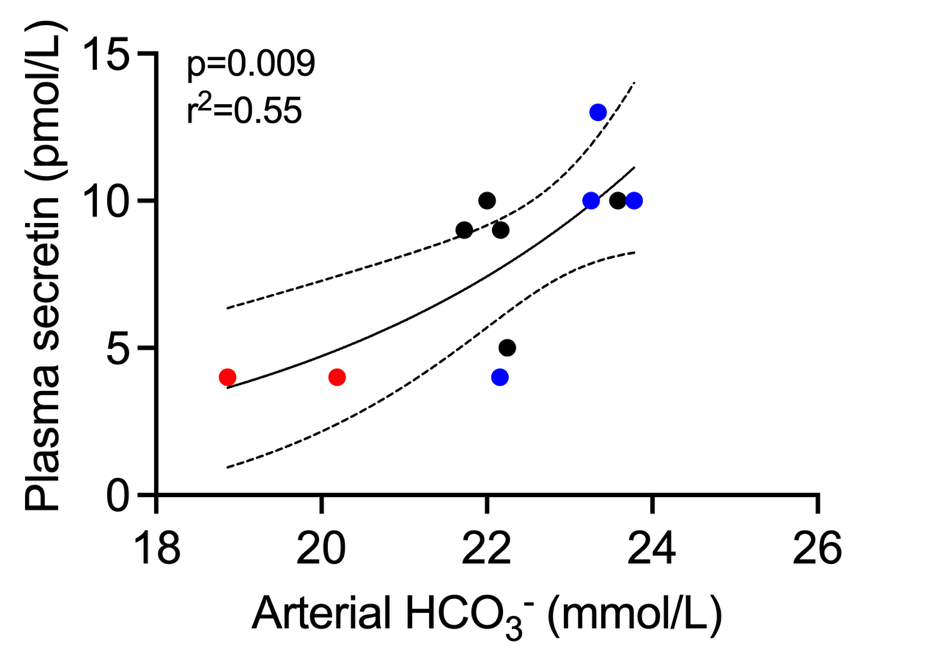

Association between arterial HCO_3_^-^ and plasma secretin in female C57Bl/6J mice exposed to control (black dots), NH_4_Cl (200 mM, red dots) or NaHCO_3_ (200 mM, blue dots) drinking water for 24 hours.

#### Figure S7

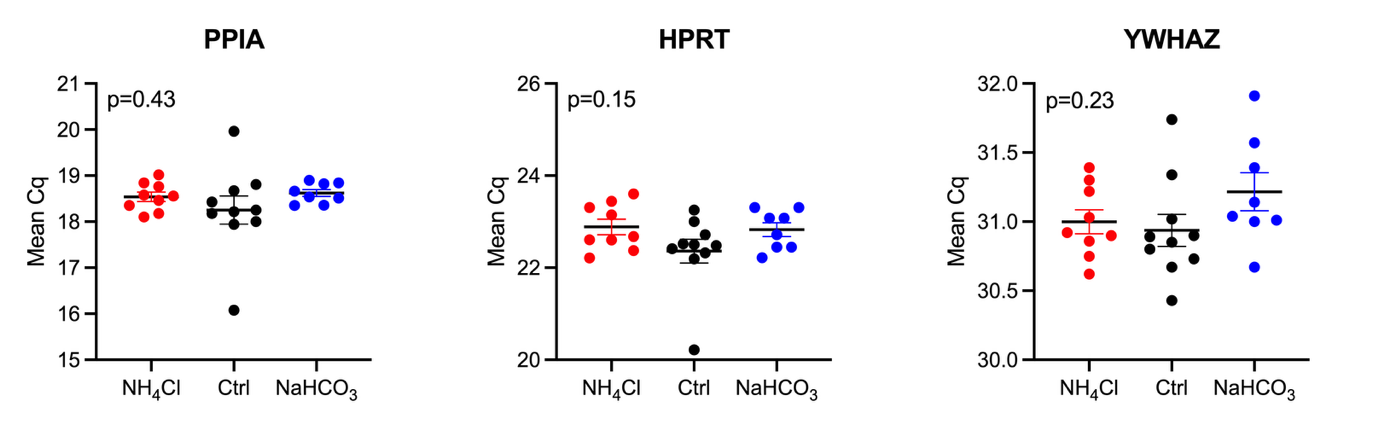

Mean Cq values of the two house-keeping genes used to normalize SCTR expression. Differences across groups were assessed by one-way ANOVA.

#### Figure S8

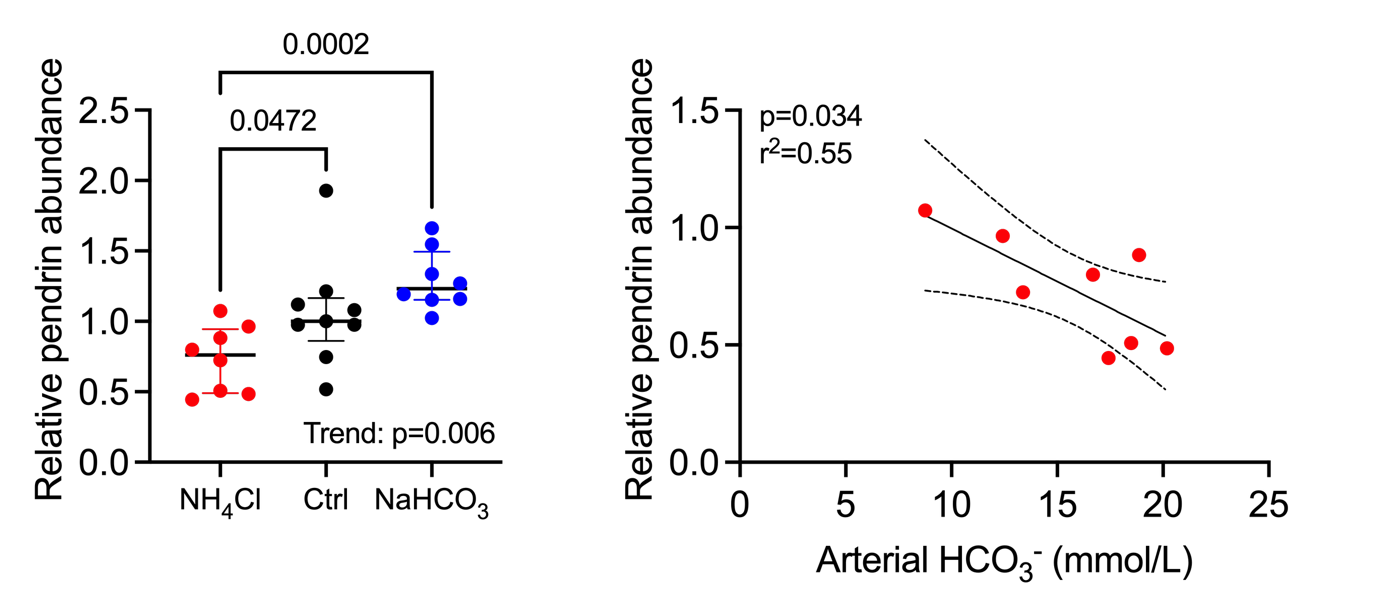

Relative pendrin protein abundance in C57Bl/6J mice exposed to control (black dots), NH_4_Cl (200 mM, red dots) or NaHCO_3_ (200 mM, blue dots) drinking water for 24 hours (left panel) and association between arterial HCO_3_^-^ and pendrin protein abundance in acid-loaded (200 mM NH_4_Cl) mice (right panel). Pendrin protein abundance was normalized to total protein loading using post chemiluminescent Coomassie staining and expressed relative to the mean of the control group (black dots).

#### Figure S9

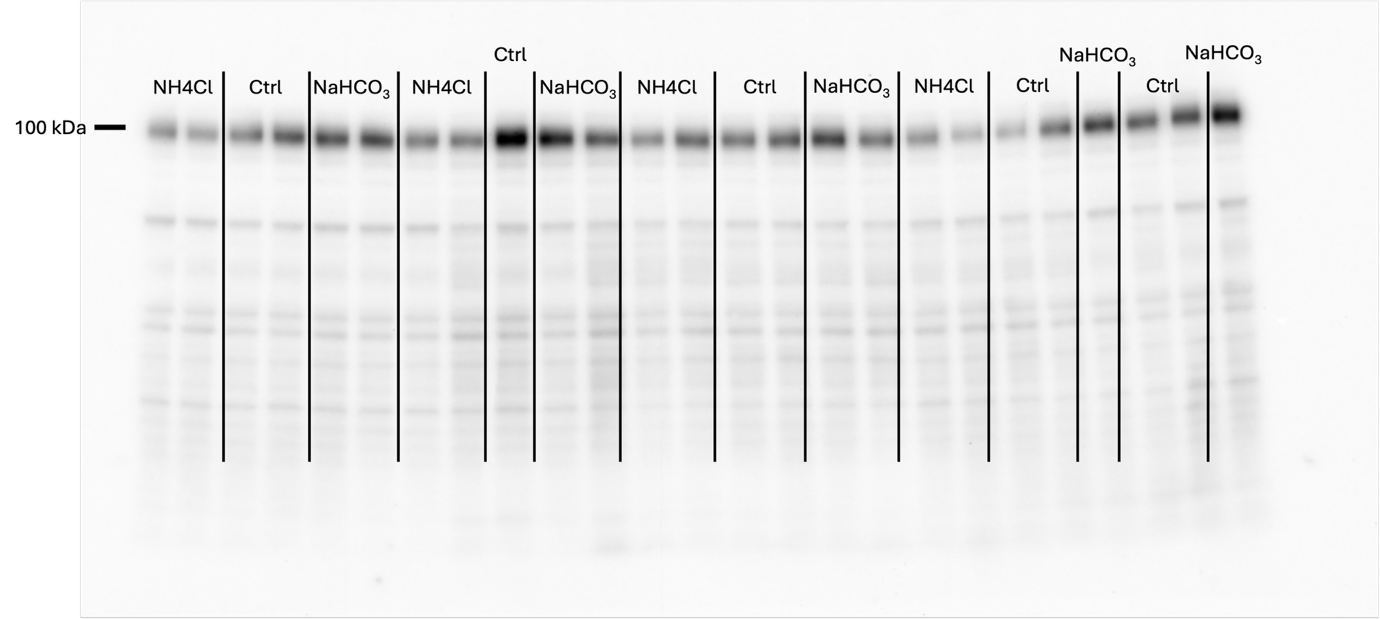

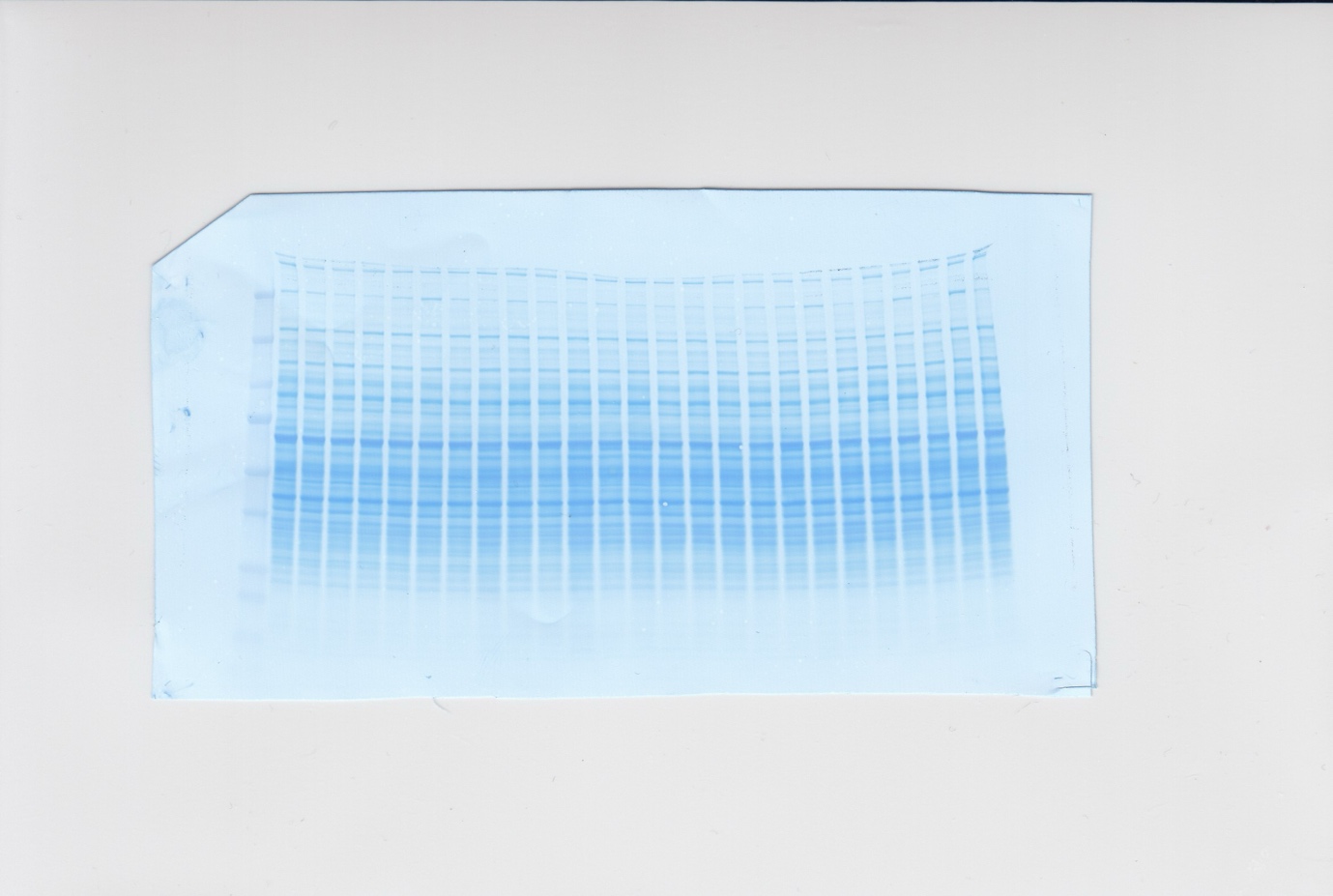

Uncropped full-length pendrin blot and corresponding Coomassie staining of the membrane of kidney lysates from C57Bl/6J mice exposed to control, NH_4_Cl (200 mM) or NaHCO_3_ (200 mM) drinking water for 24 hours.

1.	Vitzthum H, Koch M, Eckermann L, et al. The AE4 transporter mediates kidney acid-base sensing. *Nat Commun.* 2023;14(1):3051.
